# Real-Time Imaging of Antimalarial Drug Effects on *Plasmodium falciparum*-Infected Red Blood Cells

**DOI:** 10.1101/2024.11.21.624618

**Authors:** Julia S. Wallner, Maike Bonsen, Emilie Mathis, Ana Rita Gomes, Thomas Spangenberg, Julien Guizetti

## Abstract

Malaria remains a significant global health challenge, exacerbated by emerging drug resistance, which has been recurring ever since Chloroquine resistance was reported to spread in the 1960s. Cabamiquine (M5717), a promising antimalarial candidate, targets *Plasmodium falciparum* elongation factor 2 (PfeEF2) and has demonstrated high potency and multi-stage activity. However, differences in parasite-killing kinetics require further exploration. This proof-of-concept study employs time-lapse microscopy to assess the effects of Chloroquine and Cabamiquine on *P. falciparum*-infected red blood cells. Using a fluorogenic DNA dye, we monitored morphological changes and cell death at the single-cell level. Unlike the faster-killing Chloroquine, our findings confirm a lag phase of the effect of Cabamiquine and reveal a delay before host cell lysis. This delayed cytotoxic effect underscores the cytostatic phase of the drug prior to killing and highlights its potential in combination therapies. Our novel imaging approach provides a comprehensive fingerprinting method to quantify cytostatic, cytotoxic, and cytolytic activities of antimalarial drugs, offering valuable insights into their mechanisms of action at a cellular level therefore enabling to discriminate the steps prior to parasite clearance in hosts.

## Introduction

Malaria is caused by intracellular parasites of the genus *Plasmodium* and remains a severe burden on global health. Recently, a resurgence in cases has been observed, resulting in over 249 million patients and approximately 608,000 malaria-related deaths (WHO, 2022). One major challenge in combating the disease is the emergence of resistance to antimalarial drugs (Haldar et al, 2018). A notable example is Chloroquine, a hemozoin formation inhibitor,, widely used from the 1940s until the 1960s, before widespread drug resistance emerged in Africa (Tse *et al*, 2019). More recently, treatment failures with the frontline antimalarial Artemisinin have highlighted the urgent need for novel drugs (Dondorp, 2010; Ariey *et al*, 2014; Dondorp *et al*, 2009; Ashley *et al*, 2014).

A key strategy to reduce the risk of drug resistance is the implementation of combination therapy. Identifying new partner drugs with novel modes of action is therefore a priority in malaria research. One promising candidate is Cabamiquine (M5717), which targets the elongation factor 2 (PfeEF2) of *Plasmodium falciparum*, essential for protein synthesis (Baragana *et al*, 2015). Cabamiquine’s long half-life, multi-stage antiplasmodial activity, and high potency make it an attractive drug, which has successfully completed Phase 1 safety trials, ClinicalTrials.gov (NCT03261401), (McCarthy *et al*, 2021; Plas *et al*, 2023; Stadler *et al*, 2023). It is currently undergoing Phase 2 clinical trials as a combination treatment with Pyronaridine, ClinicalTrials.gov (NCT05974267), (Rottmann *et al*, 2020; Fontinha *et al*, 2022).

Despite its potency, studies have seen a delay in parasite-killing using parasite reduction rate assays, which detects parasite resurgence after wash out of the drug within 24-48 h (Baragana *et al*, 2015; Sanz *et al*, 2012). Other studies have also noted a delayed clearance (Parkyn Schneider *et al*, 2023), yet the kinetics with which Cabamiquine arrests, kills, or lyses parasites in the blood have not been fully explored.

This issue reflects a broader challenge in antiparasitic drug testing. Common metrics like parasite reduction ratio (PRR) and parasite clearance time (PCT) do not fully capture the impact on the parasite population (Khoury *et al*, 2020) i.e. the discrimination between the action of the drug, e.g. protein synthesis arrest, the ensuing parasite death (killing) and at last the lysis of the infected red blood cell (iRBC). Additionally, metabolic tests, such as lactate dehydrogenase activity assays, cannot readily differentiate between cytostatic and cytotoxic effects.

To better discriminate between those effects of antimalarial drugs in vitro, we leverage live cell imaging of *Plasmodium*-infected red blood cells, a technique that has not yet been extensively explored for that purpose. Here, we introduce a proof-of-concept approach using time-lapse microscopy to quantitatively assess morphological changes and the timing of death in individual parasites. With the appropriate markers this approach offers the potential to quantify cytostatic, killing and lysing activity of a drug in one in vitro assay at single cell level. Our data using fluorogenic DNA dyes as a marker for cell permeability confirms that Cabamiquine kills parasites after a lag phase that is not seen for Chloroquine, and reliably halts cell cycle progression within 6h. Both drugs display a delay between death-associated changes of dye permeability and lysis of the host cells, which is more pronounced for Chloroquine.

## Results and Discussion

### Long term imaging protocol enables comparison between IDC event dynamics

To assess the impact of 5 nM Cabamiquine (CBQ) on individual *P. falciparum* blood stage parasites, we acquired time lapse imaging data of the 3D7 wild type strain using an Airyscan confocal microscope in low phototoxicity mode under physiological incubation conditions. As a benchmark for fast-acting and fast-clearing drugs, we used Chloroquine (CQ) at 25 nM also corresponding to about five times the IC50 (Tremblay *et al*, 2023; Kreidenweiss *et al*, 2008). To visualize parasite nuclei in addition to the infected red blood cell (iRBC) morphology we added a next-generation infrared fluorogenic live cell dye 5-SiR-Hoechst about 4 h before the start of the movie (Fig. 1A) (BuceviČius *et al*, 2019; Wenz *et al*, 2022; Klaus *et al*, 2022). Drug or control were added to the imaging dish about 1 h before start of acquisition.

**Figure 1.**
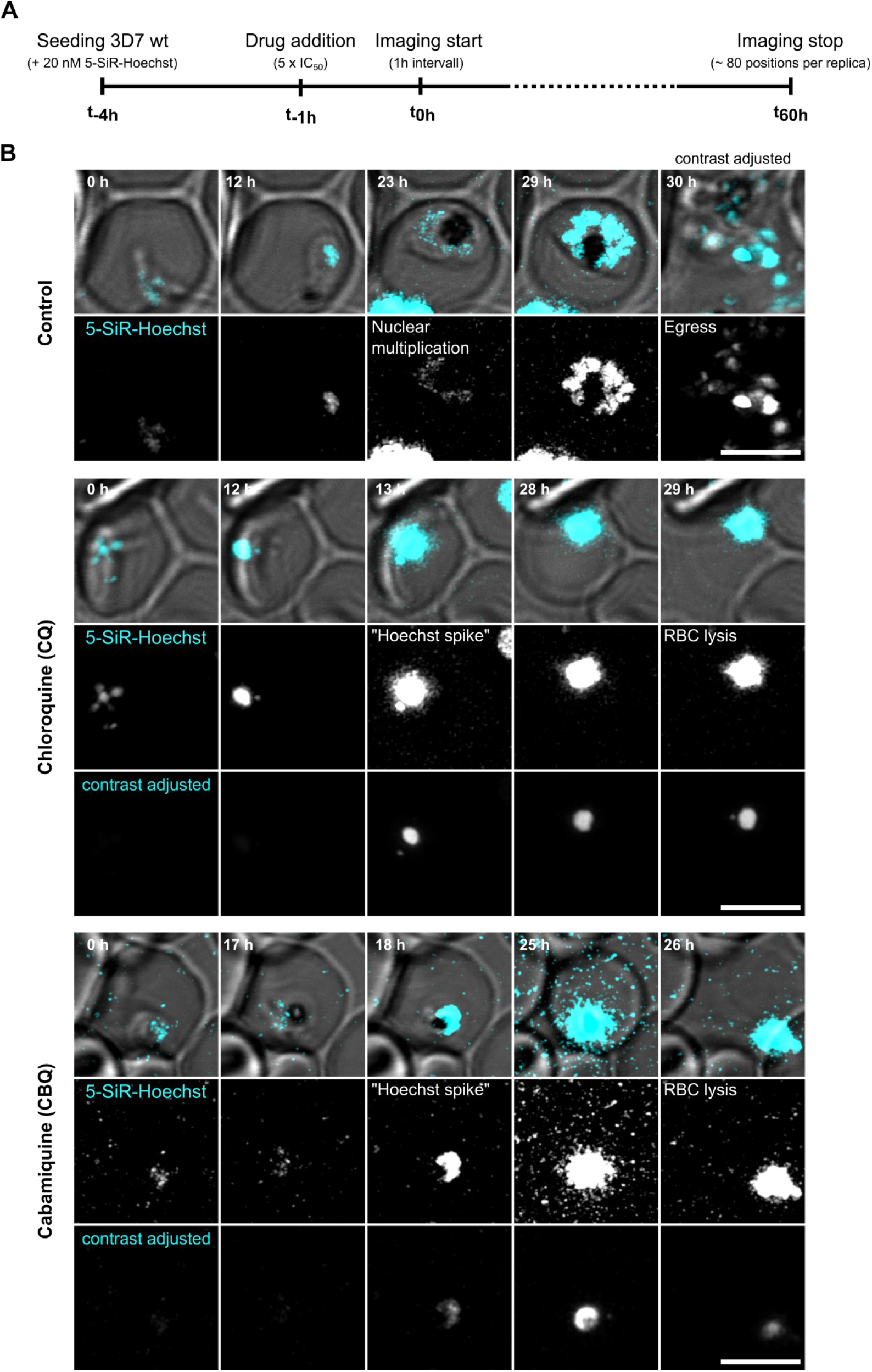
Long-term time-lapse microscopy of drug treated P. falciparum-infected red blood cells. **A** Schematic timeline of parasite seeding into imaging dishes, labeling, treatment conditions and imaging protocol **B** Representative stills from time lapse movies acquired of 3D7 parasites labeled with 20 nM 5-SiR-Hoechst (cyan) on Zeiss LSM 900 Airyscan that were either untreated, or to which 5 nM Cabamiquine, or 25 nM Chloroquine were added. Transmission channel (grey). Due to the very strong fluctuation of the signal during Hoechst spike and merozoite egress the contrast adjusted channels have been provided for adequate visibility. Scale bars, 5 μm.

Parasites of different stages were randomly selected from an asynchronous population, largely omitting schizont stages as they had already begun dividing their nuclei and drug-treated wells and imaged for 60 h at 1 h time intervals. In most control parasites we detected nuclear multiplication followed by egress from the iRBC concluding the intraerythrocytic development cycle (IDC) (Fig. 1B). Drug-treated parasites, however, failed to undergo nuclear multiplication or egress, but rather showed a substantial and rapid increase in Hoechst fluorescence intensity followed with some delay by RBC lysis in most cases. We manually scored this event when noticing a >4-fold increase of maximum Hoechst signal intensity within 1-2 hours and designated it as “Hoechst spike”. Despite being more challenging to quantify in this assay, we neither detected substantial parasite nor hemozoin growth under drug treatment.

SiR-Hoechst dyes are known to be actively pumped out of cells, which has led to the suggestion to add verapamil, an efflux pump inhibitor, for improved staining (Lukinavicius *et al*, 2015, 2014). Due to the *P. falciparum* being highly sensitive to Verapamil, an inhibitor of the drug efflux pump, we have not exploited that option, particularly in the context of live cell imaging (Martin *et al*, 1987). Differential permeability to Hoechst dyes has also been confirmed in *P. falciparum* (Sturm *et al*, 2009). It is further important to note that the permeability of the dye changes as the iRBC matures leading to a gradual increase in staining in late schizonts and particularly once the merozoites egress from the RBC (Fig. 1B). The sudden and drastic increase of 5-SiR-Hoechst fluorescence intensity prior to IDC completion, however, can be interpreted as a loss metabolic activity. The lack of growth after 5-SiR-Hoechst influx, the association with host cell lysis, and the substantially higher occurrence in drug-treated parasites suggest that it can be used as a proxy for parasite death or at least cytotoxic activity. Premature lysis of the iRBC can be considered as definitive parasite death marker in this obligate intracellular life cycle stage (Fig. 1B).

We also experimented with various Mitotracker dyes, which have been successfully used as endpoint markers (Moon *et al*, 2013; Jogdand *et al*, 2012; Dembele *et al*, 2017), and confirmed that some show a significant loss of fluorescence intensity upon parasite death (data not shown). However, they did not allow normal IDC progression beyond a few hours once the cells were labeled precluding their use for long term live cell imaging. Considering the scarcity of robust live cell compatible live/dead stains for *Plasmodium* (Linzke *et al*, 2020), the spike in 5-SiR-Hoechst single might presents an interesting option to track parasite killing or at least cytostatic activity at the single cell level in real-time. The advantage of using fluorescent dyes is the applicability to any parasite strain without the need for genetic manipulation.

This long-term imaging protocol allows for a detailed comparison of drug effects on the IDC, providing new insights into the dynamics of parasite response to treatment.

### Drug treatment halts normal progression through the IDC

For our first quantitative analysis of parasite death-associated markers, we focused on single infected ring and trophozoite stages. This allowed us to monitor the occurrence of nuclear multiplication and egress and we found that most control cells move through the IDC normally while those events were virtually never detected in drug-treated parasites (Fig. 2A). Time-lapse data allowed us to generate a cumulative histogram of egress events of the control cells (Fig. 2B). First egress events were observed after around 10 h and plateaued at around 50 h. Considering an about 48 h IDC, this matches well with the distribution of selected cells that ranged from very early rings, which could be as early as 0-2 hours post invasion (hpi), to late trophozoites up to 32 hpi. This indicates that IDC progression was not significantly disturbed by 5-SiR-Hoechst as published previously (Wenz *et al*, 2022). Even though about 24 % of control parasites fail to egress we have shown repeatedly that long term live cell imaging of *P. falciparum* is feasible (Stürmer *et al*, 2023; Wenz *et al*, 2022; Simon *et al*, 2021; Klaus *et al*, 2022). We speculate that phototoxicity contributes to RBC lysis thereby causing death of some parasites, which is supported by the notion that percentage of iRBC lysis in controls varies between blood donations (data not shown). It is, however, worth noting that when tracking IDC progression of a bulk parasite population, e.g. by Giemsa smear, we might not notice a small drop in parasitemia prior to the next reinvasion cycle. Taken together, these data confirm that our assay, despite some RBC lysis across samples, can effectively monitor normal IDC progression in real-time, paving the way for more precise and dynamic assessments of antimalarial efficacy.

**Figure 2.**
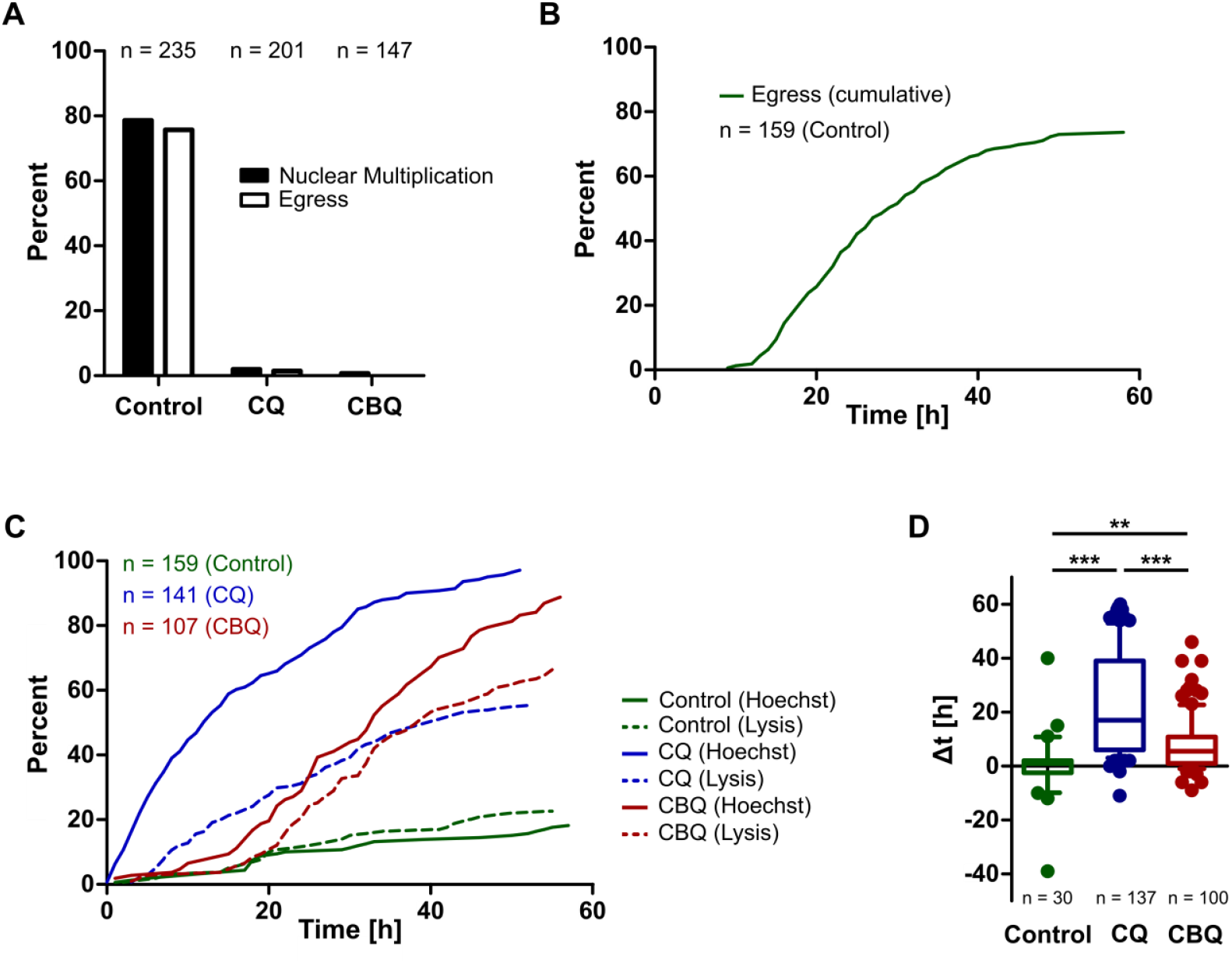
Time course of death-associated events displays different kinetics in early- and mid-stage parasites treated with drugs. **A** Percentage of parasites showing nuclear multiplication or egress within the 60-h imaging time course **B** Cumulative histogram of egress events in control parasites over time. Data was pooled from three independent replicas for each condition. **C** Cumulative histogram of “Hoechst spike” and iRBC lysis events in control and drug treated blood stage parasites over time. **D** Distribution of observed delay between “Hoechst spike” and RBC lysis for control and drug treated conditions. Statistically significant differences were assessed using Kruskal-Wallis test and Dunn’s post test (** indicates p-value < 0.01, *** indicates p < 0.001). Data was pooled from three independent replicas for each condition.

### Cabamiquine displays lag in killing kinetics and host cell lysis

Since none of the drug treated parasites displayed normal IDC progression, we quantified occurrence of “Hoechst spikes” and iRBC lysis (Fig. 2C). Within 60 h, a Hoechst spike was observed in 18 % of control cells while drug treatment caused an increase to 97 % and 89 % for Chloroquine and Cabamiquine, respectively. The Hoechst spike was usually followed by iRBC lysis in 23 % for control and 55 % and 66 % for Chloroquine and Cabamiquine, respectively, within the first 60 h. Prolonging the image acquisition time could allow presenting the full kinetics in the future. A prominent difference between both drugs was the significantly faster kinetics of parasite death, i.e. Hoechst spike, caused by Chloroquine leading to more than half of parasites being affected within 12 h while for Cabamiquine it took 33 h. While the slope in the “killing phase” was similar, Cabamiquine displayed a significant lag phase of about 15 h before onset of parasite death. Importantly, within this lag phase Cabamiquine had cytostatic action as many of controls cells would have undergone nuclear multiplication within this time. These findings align with Chloroquine being a fast-killing drug, while Cabamiquine has previously shown delayed parasite killing (Baragana *et al*, 2015). Another notable feature revealed using this type of imaging-based analysis is the delay between Hoechst spike and iRBC lysis (Fig. 2D). While some variability can be observed in control cells, and even occasionally iRBC lysis preceding the Hoechst spike, both events remain closely linked. In drug treated cells a significant delay of about 22.5 and 7.7 h for Chloroquine and Cabamiquine, respectively, can be observed between both events. These differences in phenotypes underline the different modes of action for both drugs.

### Cabamiquine has slower cytotoxic kinetics, but similar cytostatic effects compared to Chloroquine on schizont stage parasites

A critical question that remains is whether despite delayed killing the progression through the IDC is halted immediately upon drug treatment or in other words whether the drugs have a rapid cytostatic effect. To test this, we specifically treated schizont stage parasites that were approaching the end of their IDC and already had multiple nuclei and imaged them as above for 18 h. A rapidly acting drug would likely prevent egress of these late-stage parasites, while control parasites would complete their cycle within this time. Indeed, 93 % of control schizont parasites egressed within 11 h as expected for a normal cell cycle progression (Fig. 3A). Of the treated parasites, however, 13 % of the Chloroquine-treated and 11 % of the Cabamiquine-treated ones achieved egress. These events occurred only within 6 h after treatment and in schizonts that were already very close to the end of their cycle. Overall schizonts were more susceptible to drug treatment than early- or mid-stage parasites (Fig. 2C) with half of them showing a “Hoechst spike” after about 3 h of exposure for Chloroquine and 11 h for Cabamiquine (Fig. 3B). Otherwise, the kinetics qualitatively reproduced the previous observation that Chloroquine causes more rapid death and a more pronounced delay between Hoechst spike and RBC lysis. Yet, it seems that despite this difference in killing kinetics Cabamiquine is similarly effective in arresting IDC progression of parasites than Chloroquine.

**Figure 3.**
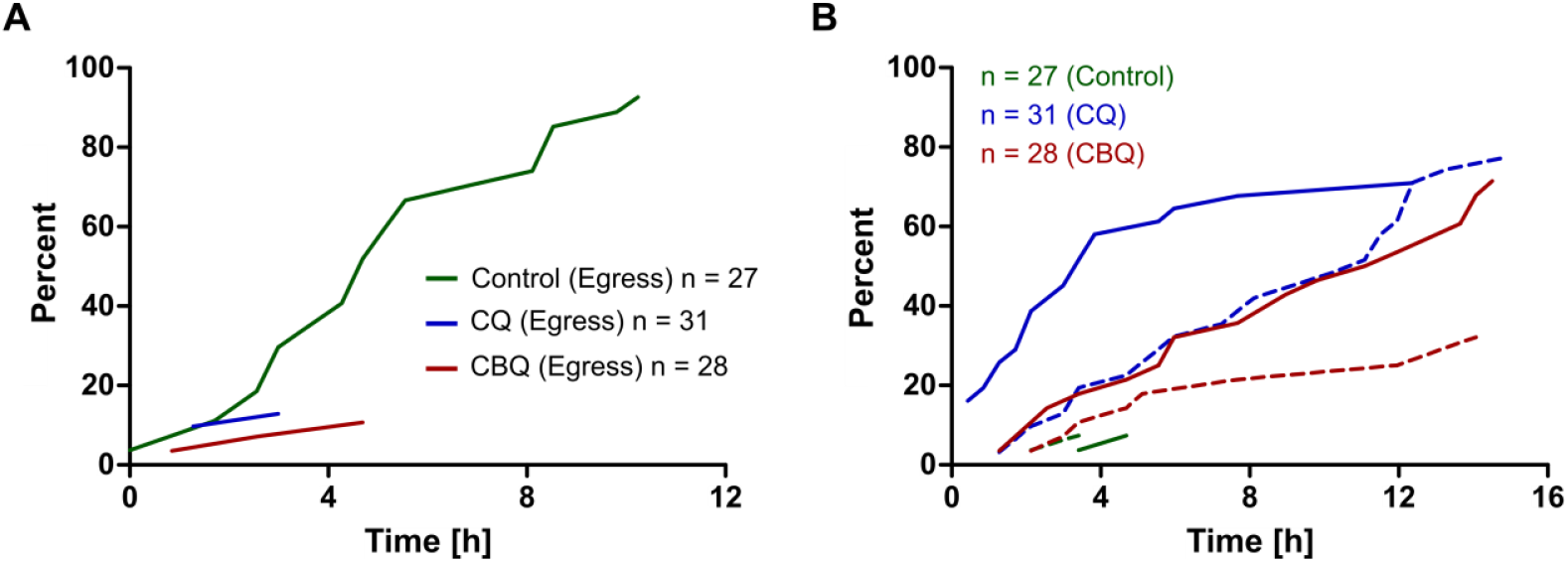
Time course of IDC progression and death-associated events in drug treated schizonts. **A** Cumulative histogram of egress events in control parasites over time. Data was pooled from two independent replicas for each condition. **B** Cumulative histogram of “Hoechst spike” (solid lines) and iRBC lysis events (dotted lines) in control and drug treated blood stage parasites over time.

## Conclusion

Our proof-of-concept study demonstrates the value of long-term live-cell imaging to assess the dynamics of drug action on *Plasmodium falciparum* at the single-cell level. This was exemplified with two antimalarial drugs with distinct modes of action i.e. Chloroquine and Cabamiquine.

Chloroquine, a hemozoin formation inhibitor was able to rapidly block parasite egress within 6h and half of parasites were killed within 12 h after exposure. The iRBC lysis followed 22.5h thereafter. Cabamiquine, a protein synthesis inhibitor was able to rapidly block parasite egress within 6h and half of parasites being killed within 33 h after exposure. The iRBC lysis in 7.7h therafter.

Overall both drugs displayed fast acting profiles to arrest parasites in their development. While the slope in the “killing phase” were similar, Cabamiquine displayed a significant lag phase of about 15 h before onset of parasite death compared to Chloroquine.. Discriminating between those effects is enabled by the assay presented here.. We lay the groundwork to investigating differences in drug susceptibility in more detail and can inform models that predict parasite growth arrest, killing and lysis for emerging candidate drugs. Future studies should aim to extend the imaging duration and incorporate additional markers to refine our understanding of drug-induced changes in parasite physiology to optimize malaria treatment regimes.

## Materials and Methods

### Parasite culturing

Parasites were cultured at 5% CO_2_ and 3% O_2_ in a humidity-controlled environment and maintained at a haematocrit of about 2% in a 6 well dish, 50 μL blood and 2 mL complete RPMI (cRPMI) were addedd per well. cRPMI medium is made with 500 mL RPMI1640 (Thermo, 61870-010, GlutaMAX, Phenol, no HEPES), 10 mL 100x Hypoxanthin (CCPro Z-41-M 100 mL) [0.2 mM], 125 μL Gentamycin (Roth HN09.1 20 mL, 50 mgmL−1 [12.5 μgmL−1], 25 mL 10% Albumax (Life 11021-029, 25 g in 500 mL RPMI aliquots at −20°C) [0.5%], 5 mL HEPES 2.5 M stock, pH 7.3 [25 mM] (Roth 5.95g in 10 mL H_2_O). Parasitaemia was maintained at ≤ 5% and monitored via Giemsa stain.

### Parasite seeding and drug treatment

Parasite cultures with a high parasitaemia of about 5% (haematocrit of 2%) are used for seeding. For a surface treated quad dish (ibidi, 80416) around 1 mL of a culture is used and needs to be spun down for 30 s at 1000 g. The pellet is then washed with 1 mL sterile PBS and spun down again, followed by the addition of 1 mL sterile PBS. 200 μL of this cell suspension is added to each well of the quad dish and incubated for 10 min. The wells are then gently washed with 200 μL of sterile PBS until there is an even monolayer on the bottom of each well. 500 μL of cRPMI with 20 nM 5-SiR-Hoechst is added to the monolayer. The seeded quad dish then stays in the incubator for around 3h allowing the cells to take up the dye efficiently. before, Thereafter the drug is added to the imaging wells in 100 μL cRPMI to 5 nM Cabamiquine (Biozol, MCE-HY-117684) or 25 nM Chloroquine (Sigma, C6628) final concentration. For controls 100 μL prewarmed cRPMI is added instead. Immediately afterwards the dish is transferred to the microscope incubation chamber with the lid being slightly open to allow for gas exchange.

### Time-lapse imaging

For live cell imaging a Zeiss LSM 900 Airyscan 2 microscope in sensitivity mode with a 63x 1.40NA oil objective was used. The incubation chamber was heated to 37 °C and gased with 3% CO_2_ and 5% O_2_. The illumination settings for transmission channel 1 used T-DMT detector with 640 nm laser at 0.02% power and for infrared channel 2 we used the Airyscan detector with 640 nm laser at 0.2% power at a scan speed 7 resulting in a pixel dwell time of 2.32 μs. Image settings are 56 nm pixel size, 454×454 pixels per image. The z-stacks were 6 μm with 13 slices at an interval of 500nm. Multiple infected RBCs were excluded from analysis. Images were acquired every hour for 60 h total. The image data was enhanced by Airyscan processing. The data were pooled from three independent replicas for each condition.

### Image analysis

Movies were manually annotated after drift correction and contrast adjustment using Fiji/ Image J (Schindelin *et al*, 2012). iRBCs without any clear hemozoin pigment were classified as ring stages. Small hemozoin containing parasites were classified as early trophozoites and larger parasites with prominent hemozoin as late trophozoites. Schizonts were classified by checking if more than one distinct nucleus could be detected. Nuclear multiplication was scored when multiple clearly distinguishable nuclei appeared. Egress was scored when iRBCs burst after formation of distinct merozoites. This could be differentiated from RBC lysis, which occurs before parasites have reached the late schizont stage. Hoechst spike was scored when the maximal fluorescence intensity within a parasite nucleus increased by more than 4-fold within one or two time frames.

## Author Contributions

Experiments were planned and designed by J.G with help from T.S. J.S.W and J.G carried out the experiments with the help of M.B and E.M, who is supervised by A.R.G. Data analysis was performed by J.S.W with the help J.G. Figures were generated by J.G. The manuscript was written by J.G with input from J.S.W and T.S.

## Acknowledgments

We thank: The Infectious Diseases Imaging Platform for imaging support (idip-heidelberg.org). Cecilia Sanchez for providing Chloroquine. Claudia Demarta-Gatsi for initial discussions of the project.

## Data and Material Availability Statement

All data is provided in the manuscript. Materials can be provided upon request to the corresponding author.

## Declaration of Generative AI and AI-assisted technologies in the writing process

During the preparation of this work the authors used ChatGPT 4 to revise some of the text for errors and clarity. After using this, the authors reviewed and edited the content as needed and take full responsibility for the content of the publication.

## References

Ariey F, Witkowski B, Amaratunga C, Beghain J, Langlois A-C, Khim N, Kim S, Duru V, Bouchier C, Ma L, et al (2014) A molecular marker of artemisinin-resistant Plasmodium falciparum malaria. Nature 505: 50–55

Ashley EA, Dhorda M, Fairhurst RM, Amaratunga C, Lim P, Suon S, Sreng S, Anderson JM, Mao S, Sam B, et al (2014) Spread of artemisinin resistance in Plasmodium falciparum malaria. N Engl J Med 371: 411–423

Baragana B, Hallyburton I, Lee MC, Norcross NR, Grimaldi R, Otto TD, Proto WR, Blagborough AM, Meister S, Wirjanata G, et al (2015) A novel multiple-stage antimalarial agent that inhibits protein synthesis. Nature 522: 315–320

BuceviČius J, Keller-Findeisen J, Gilat T, Hell SW & LukinaviČius G (2019) Rhodamine-Hoechst positional isomers for highly efficient staining of heterochromatin. Chemical Science 12: 1962– 1970

Dembele L, Ang X, Chavchich M, Bonamy GMC, Selva JJ, Lim MY-X, Bodenreider C, Yeung BKS, Nosten F, Russell BM, et al (2017) The Plasmodium PI(4)K inhibitor KDU691 selectively inhibits dihydroartemisinin-pretreated Plasmodium falciparum ring-stage parasites. Sci Rep 7: 2325

Dondorp AM (2010) Artemisinin Resistance in Plasmodium Falciparum. Nature Reviews Microbiology 8: 272–80

Dondorp AM, Nosten F, Yi P, Das D, Phyo AP, Tarning J, Lwin KM, Ariey F, Hanpithakpong W, Lee SJ, et al (2009) Artemisinin resistance in Plasmodium falciparum malaria. N Engl J Med 361: 455–467

Fontinha D, Arez F, Gal IR, Nogueira G, Moita D, Baeurle THH, Brito C, Spangenberg T, Alves PM & Prudêncio M (2022) Pre-erythrocytic Activity of M5717 in Monotherapy and Combination in Preclinical Plasmodium Infection Models. ACS Infectious Diseases

Haldar K, Bhattacharjee S & Safeukui I (2018) Drug resistance in Plasmodium. Nature Reviews Microbiology

Jogdand PS, Singh SK, Christiansen M, Dziegiel MH, Singh S & Theisen M (2012) Flow cytometric readout based on Mitotracker Red CMXRos staining of live asexual blood stage malarial parasites reliably assesses antibody dependent cellular inhibition. Malar J 11: 235

Khoury DS, Zaloumis SG, Grigg MJ, Haque A & Davenport MP (2020) Malaria Parasite Clearance: What Are We Really Measuring? Trends in Parasitology 36: 413–426

Klaus S, Binder P, Kim J, Machado M, Funaya C, Schaaf V, Klaschka D, Kudulyte A, Cyrklaff M, Laketa V, et al (2022) Asynchronous nuclear cycles in multinucleated Plasmodium falciparum facilitate rapid proliferation. Science Advances 8: 1–13

Kreidenweiss A, Kremsner PG & Mordmüller B (2008) Comprehensive study of proteasome inhibitors against Plasmodium falciparum laboratory strains and field isolates from Gabon. Malaria Journal 7: 187

Linzke M, Yan SLR, Tárnok A, Ulrich H, Groves MR & Wrenger C (2020) Live and Let Dye: Visualizing the Cellular Compartments of the Malaria Parasite Plasmodium falciparum. Cytometry Part A 97: 694–705

Lukinavicius G, Blaukopf C, Pershagen E, Schena A, Reymond L, Derivery E, Gonzalez-Gaitan M, D’Este E, Hell SW, Gerlich DW, et al (2015) SiR-Hoechst is a far-red DNA stain for live-cell nanoscopy. Nat Commun 6: 8497–8497

Lukinavicius G, Reymond L, D’Este E, Masharina A, Gottfert F, Ta H, Guther A, Fournier M, Rizzo S, Waldmann H, et al (2014) Fluorogenic probes for live-cell imaging of the cytoskeleton. Nat Methods 11: 731–733

Martin SK, Oduola AM & Milhous WK (1987) Reversal of chloroquine resistance in Plasmodium falciparum by verapamil. Science 235: 899–901

McCarthy JS, Yalkinoglu Ö, Odedra A, Webster R, Oeuvray C, Tappert A, Bezuidenhout D, Giddins MJ, Dhingra SK, Fidock DA, et al (2021) Safety, pharmacokinetics, and antimalarial activity of the novel plasmodium eukaryotic translation elongation factor 2 inhibitor M5717: a first-in-human, randomised, placebo-controlled, double-blind, single ascending dose study and volunteer infection study. The Lancet Infectious Diseases 21: 1713–1724

Moon S, Lee S, Kim H, Freitas-Junior LH, Kang M, Ayong L & Hansen MAE (2013) An image analysis algorithm for malaria parasite stage classification and viability quantification. PLoS One 8: e61812

Parkyn Schneider M, Looker O, Rebelo M, Khoury DS, Dixon MWA, Oeuvray C, Crabb BS, McCarthy J & Gilson PR (2023) The delayed bloodstream clearance of Plasmodium falciparum parasites after M5717 treatment is attributable to the inability to modify their red blood cell hosts. Front Cell Infect Microbiol 13: 1211613

Plas JL van der, Kuiper VP, Bagchus WM, Bödding M, Yalkinoglu Ö, Tappert A, Seitzinger A, Spangenberg T, Bezuidenhout D, Wilkins J, et al (2023) Causal chemoprophylactic activity of cabamiquine against Plasmodium falciparum in a controlled human malaria infection: a randomised, double-blind, placebo-controlled study in the Netherlands. The Lancet Infectious Diseases 23: 1164–1174

Rottmann M, Jonat B, Gumpp C, Dhingra SK, Giddins MJ, Yin X, Badolo L, Greco B, Fidock DA, Oeuvray C, et al (2020) Preclinical Antimalarial Combination Study of M5717, a Plasmodium falciparum Elongation Factor 2 Inhibitor, and Pyronaridine, a Hemozoin Formation Inhibitor. Antimicrob Agents Chemother 64: e02181–19

Sanz LM, Crespo B, De-Cózar C, Ding XC, Llergo JL, Burrows JN, García-Bustos JF & Gamo F-J (2012) P. falciparum In Vitro Killing Rates Allow to Discriminate between Different Antimalarial Mode-of-Action. PLOS ONE 7: e30949

Schindelin J, Arganda-Carreras I, Frise E, Kaynig V, Longair M, Pietzsch T, Preibisch S, Rueden C, Saalfeld S, Schmid B, et al (2012) Fiji: An open-source platform for biological-image analysis. Nature Methods 9: 676–682

Simon CS, Funaya C, Bauer J, Voβ Y, Machado M, Penning A, Klaschka D, Cyrklaff M, Kim J, Ganter M, et al (2021) An extended DNA-free intranuclear compartment organizes centrosome microtubules in malaria parasites. Life Science Alliance 4: e202101199–e202101199

Stadler E, Maiga M, Friedrich L, Thathy V, Demarta-Gatsi C, Dara A, Sogore F, Striepen J, Oeuvray C, Djimdé AA, et al (2023) Propensity of selecting mutant parasites for the antimalarial drug cabamiquine. Nat Commun 14: 5205

Sturm A, Graewe S, Franke-Fayard B, Retzlaff S, Bolte S, Roppenser B, Aepfelbacher M, Janse C & Heussler V (2009) Alteration of the parasite plasma membrane and the parasitophorous vacuole membrane during exo-erythrocytic development of malaria parasites. Protist 160: 51– 63

Stürmer VS, Stopper S, Binder P, Klemmer A, Lichti NP, Becker NB & Guizetti J (2023) Progeny counter mechanism in malaria parasites is linked to extracellular resources. PLOS Pathogens 19: e1011807

Tremblay T, Bergeron C, Gagnon D, Bérubé C, Voyer N, Richard D & Giguère D (2023) Squaramide Tethered Clindamycin, Chloroquine, and Mortiamide Hybrids: Design, Synthesis, and Antimalarial Activity. ACS Med Chem Lett 14: 217–222

Tse EG, Korsik M & Todd MH (2019) The past, present and future of anti-malarial medicines. Malar J 18: 93

Wenz C, Simon CS, Romão TP, Stürmer V, Machado M, Klages N, Klemmer A, Voß Y, Ganter M, Brochet M, et al (2022) An Sfi1-like centrin-interacting centriolar plaque protein affects nuclear microtubule homeostasis. bioRxiv: 2022.07.28.501831-2022.07.28.501831

WHO (2022) WHO Global, World Malaria Report 2022

